## Supplemental Figures for "Synergy between SIRT1 and SIRT6 helps recognize DNA breaks and potentiate the DNA damage response and repair"

**Supplementary Figures**

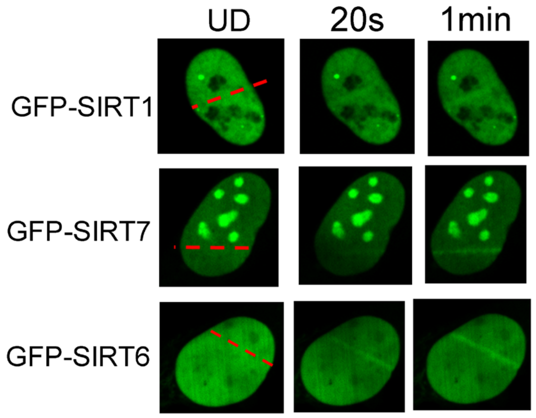


**Figure S1 (Related to Figure 1). DSB-recruitment of SIRTs**

GFP-fused SIRT1, SIRT6 and SIRT7 were introduced into MEFs. The fluorescence signal was captured 20 s and 1 min after laser-induced DNA damage.

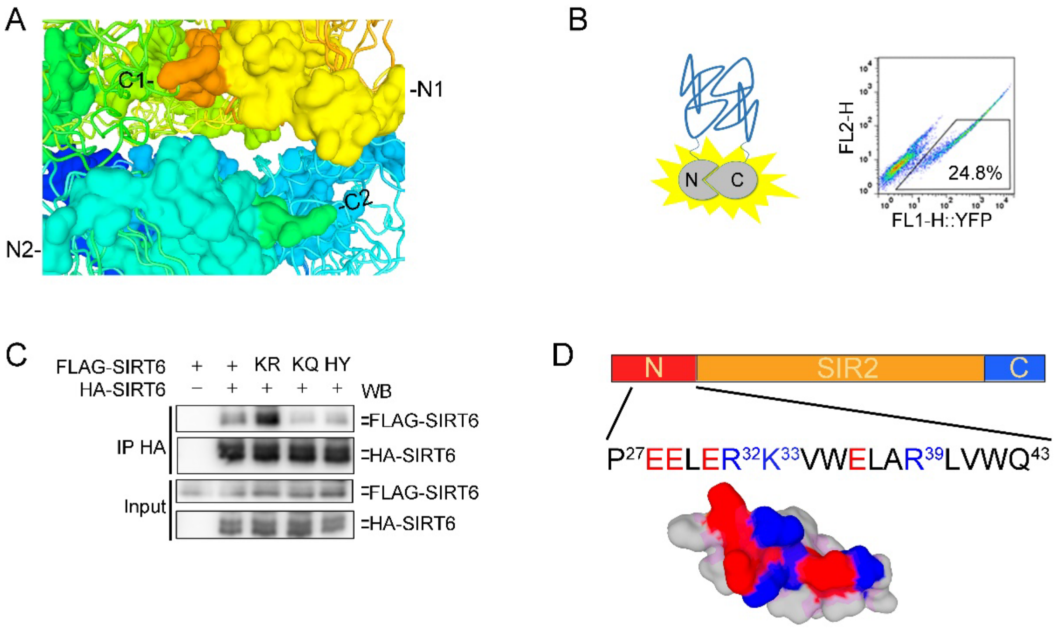


**Figure S2 (Related to Figure 2). Polymerization of SIRT6**

(**A**) Schematic of one putative DSB-binding pocket, consisting of two N-termini (yellow and aquamarine) and two C-termini (orange and lime green) of two adjacent SIRT6 hexamer molecules. (B) Left, schematic of the BiFC system; right, the yellow fluorescence detection in HEK293 cells ectopically expressing SIRT6 via the BiFC system. (**C**) Co-IP and western blot analysis of the interaction between FLAG-SIRT6 or various mutants (KR, KQ and HY) and HA-SIRT6. (**D**) Schematic of a SIRT6 N-terminal peptide showing the positively (blue) and negatively charged residues (red).

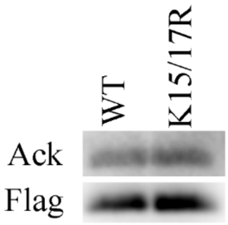


**Figure S3 (Related to Figure 2). Acetylation of SIRT6**

SIRT6 K15/17R acetylation levels, as determined by western blotting with a pan anti-acetyl lysine antibody.

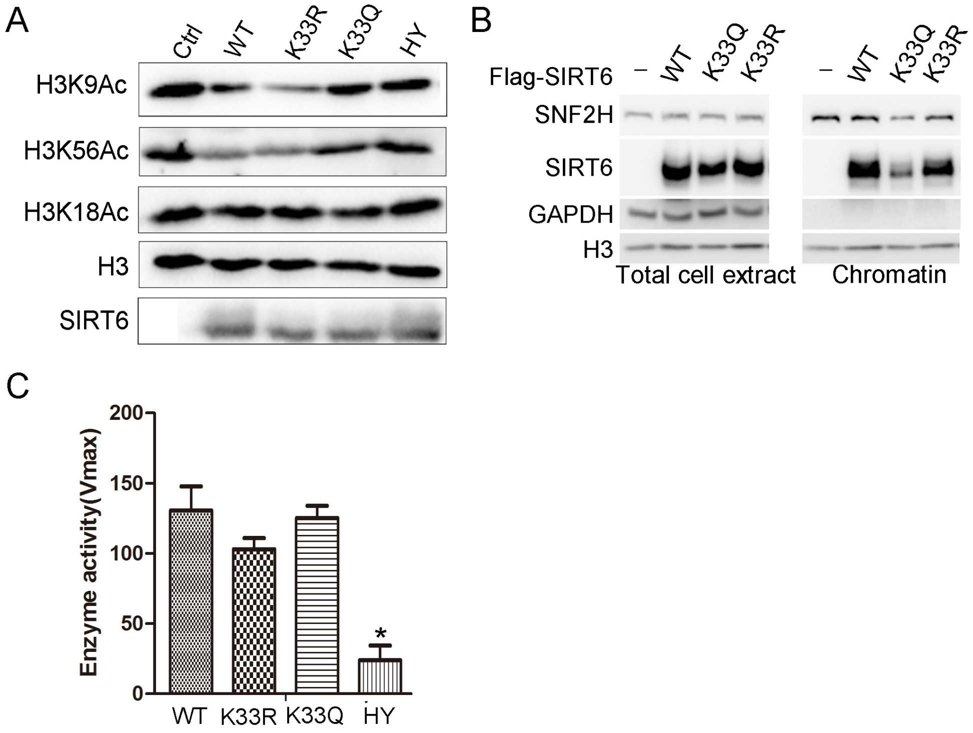


**Figure S4 (Related to Figure 2). Deacetylase activity of SIRT6**

(**A**) Western blotting analysis of the acetylation levels of histone H3 in *SIRT6* KO HEK293 cells reconstituted with SIRT6 WT or indicated mutants.

(**B**) Cell fractionation analysis to detect the chromatin enrichment of SNF2H and SIRT6 in FLAG-SIRT6, K33R, K33Q and HY reconstituted *SIRT6* KO HEK293T cells.

(**C**) Recombinant GST-SIRT6, K33R, K33Q and H133Y (HY) deacetylation activities (Vmax) *in vitro* (CycLex®). $*P < 0.05$, HY Vs WT.

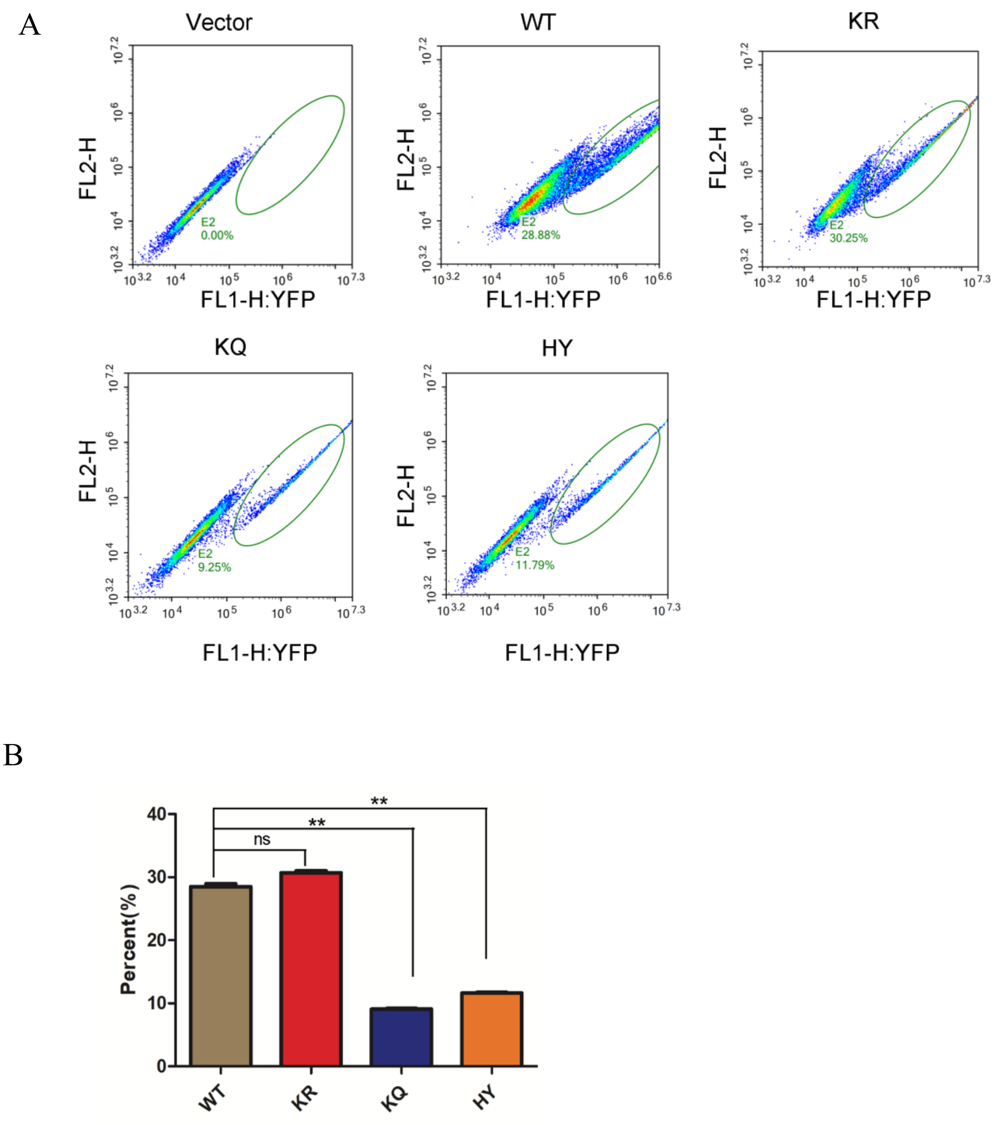


**Figure S5. Polymerization of SIRT6**

(**A**) Yellow fluorescence detection by FACS in HEK293 cells ectopically expressing vector only, SIRT6-WT, SIRT6-KR, SIRT6-KQ or SIRT6-HY via the BiFC system. Green ovals indicate yellow fluorescence positive cell population.

(**B**) The percent of yellow fluorescence positive cells from each group shown in A. $^{**}P < 0.01$, n.s., not significant.

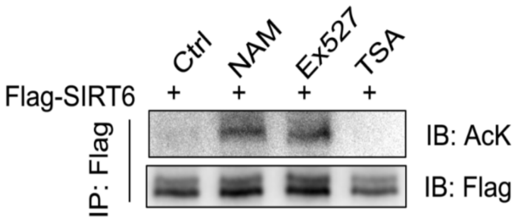


**Figure S6. Acetylation level of SIRT6**

FLAG-SIRT6 acetylation levels in the presence of NAM (5 mM), TSA (1 μM) or Ex527 (1 μM), determined by western blotting using an anti-pan-acetyl K antibody.

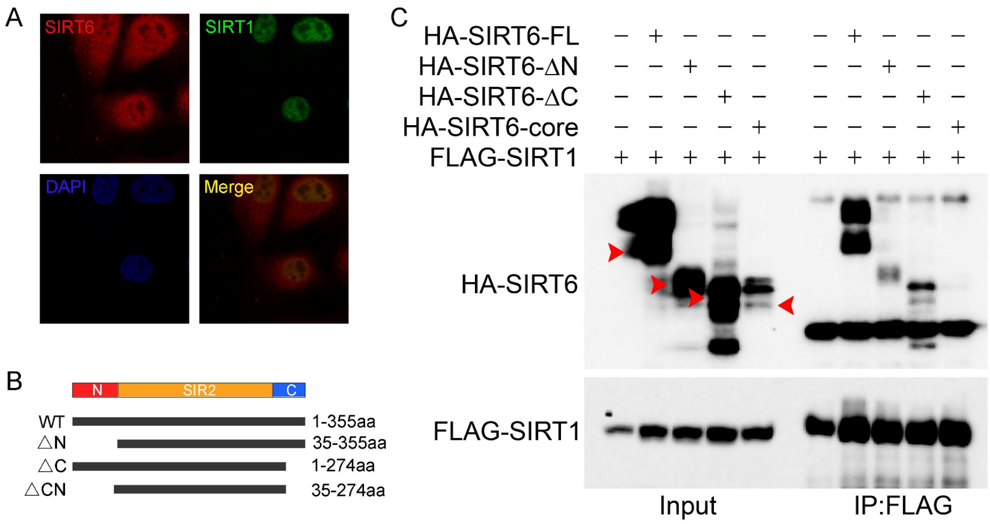


**Figure S7. SIRT1-SIRT6 interaction**

(**A**) Immunofluorescence analysis of endogenous SIRT1 (Green) and SIRT6 (Red) protein levels. Representative images are shown, captured under a confocal imaging microscope. Scale bar, 10 µm. (**B**) A schematic of the various domain-modified SIRT6 constructs. (**C**) Co-IP and western blot analysis of the interaction between SIRT1 and various domain modified SIRT6 constructs overexpressed in HEK293 cells.
